# The SUMOylation inhibitor TAK-981 (Subasumstat) triggers IFN-I-dependent activation of Natural Killer cells against Acute Myeloid Leukemias

**DOI:** 10.1101/2024.02.19.580882

**Authors:** Rawan Hallal, Marion De Toledo, Denis Tempé, Sara Zemiti, Loïs Coënon, Delphine Gitenay, Simon George, Sarah Bonnet, Ludovic Gabellier, Guillaume Cartron, Mireia Pelegrin, Martin Villalba, Guillaume Bossis

## Abstract

Natural Killer (NK) cells play a pivotal role in mounting an anti-cancer immune response. Patients with diminished NK cells number and activity face less favorable prognosis. Promising therapeutic strategies include the adoptive transfer of NK cells or the reactivation of patients’ own NK cells. TAK-981, a first-in-class inhibitor of SUMOylation undergoing phase I/II clinical trials for cancer, is emerging as an immunomodulatory drug. Here, we demonstrate that TAK-981 activates NK cells from healthy donors and patients with Acute Myeloid Leukemia (AML), a cancer with very poor prognosis. TAK-981 heightens their degranulation capacity, secretion of inflammatory cytokines (IFN-γ, TNF-α, FasL), and cytotoxicity against AML cells. *In vivo*, TAK-981 also enhances the anti-leukemic activity of *ex-vivo* expanded human NK cells. At the molecular level, TAK-981 first induces *IFNB1* gene in NK cells, leading to the secretion of type I Interferon (IFN-I), which binds to the Interferon receptor IFNAR. This induces Interferon-Stimulated Genes (ISG) and activates NK cells *in vitro* and *in vivo*. Finally, TAK-981 stimulates IFN-I secretion by monocytes, which contributes to the activation of NK cells *in trans*. Altogether, targeting SUMOylation could be a promising strategy to reactivate AML patients’ NK cells and enhance the efficiency of NK cells-based therapies.

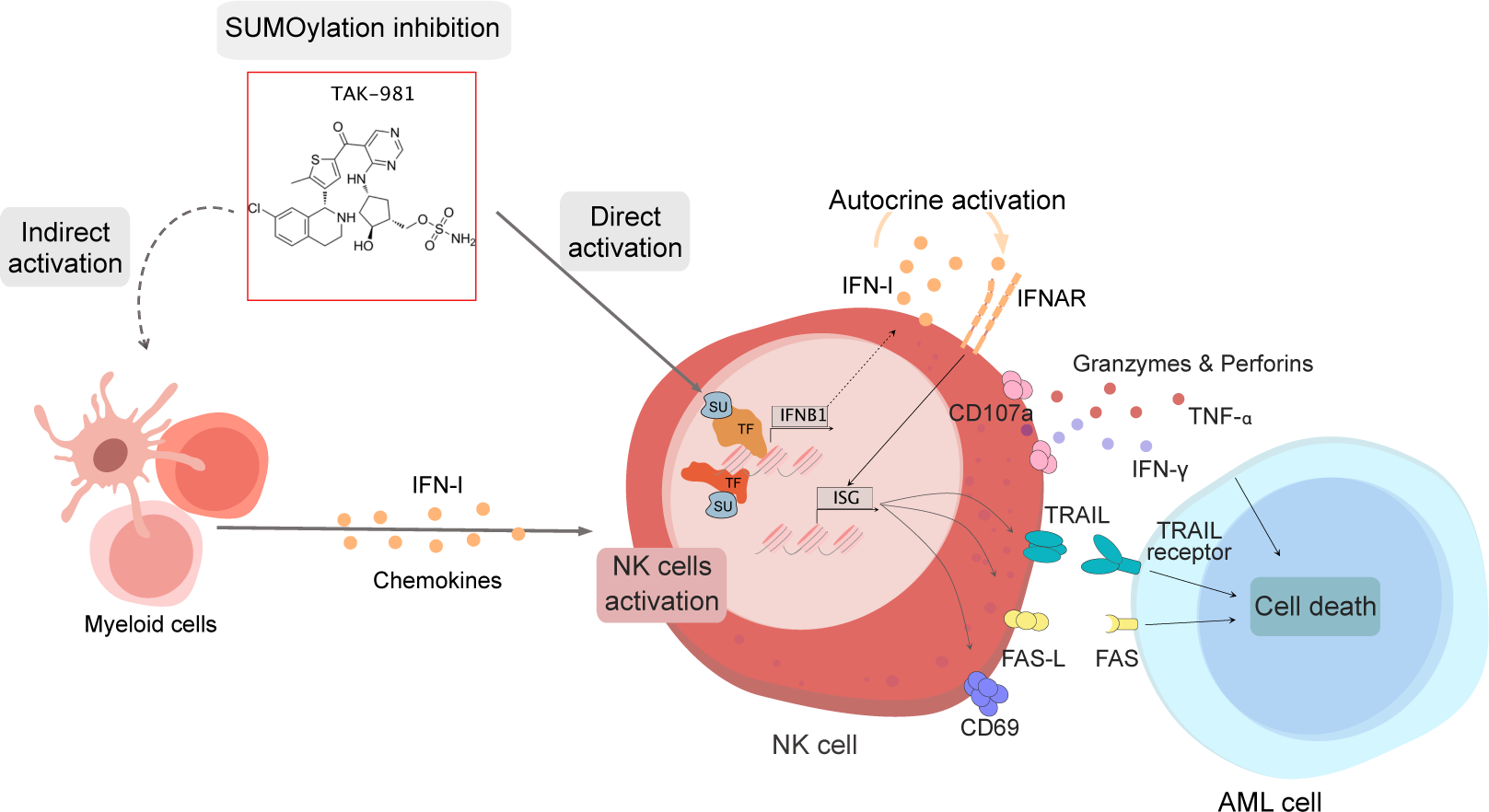

## Introduction

Natural Killer (NK) cells are lymphocytes of the innate immune system that play a crucial role in the surveillance of cancer cells. This is notably the case in Acute Myeloid Leukemias (AML), a group of aggressive myeloproliferative malignancies with poor prognosis. High numbers of circulating NK cells and higher NK cells cytotoxicity correlate with better prognosis in AML patients (1). However, frequent deficiencies in NK cells number and activity facilitate AML cells escape from immune surveillance (2). NK cells cytotoxicity indeed relies on a delicate balance between the stimulation of their activating and inhibitory receptors. NK cells from AML patients often present an increased expression of the inhibitory receptors compared to activation receptors, leading to a defective cytotoxic activity and inflammatory cytokines secretion (3). In addition, AML cells develop ways to escape NK cells by either upregulating inhibitory ligands such as the class II molecules of the histocompatibility complex (MHC-II), expressing immune-check-point markers, or shedding activating ligands from their surface (4). NK cells have also been explored as a potential cellular therapy for AML. This includes adoptive transfer of *ex vivo* expanded NK cells and chimeric antigen receptor-modified NK cells (CAR-NK). One of the main interests of allogenic NK cells transplantation lies in their potential to eliminate tumor cells without causing graft-versus-host disease (GVHD). However, new strategies are required to enhance their activation and thus strengthen their anti-tumoral activity (5).

SUMOylation is a post-translational modification, which consists in the covalent conjugation of SUMO-1, -2 or -3 on lysines of thousands of proteins to control their function and fate. One major role of SUMOylation is the control of gene expression through the modification of various transcription factors and co-regulators (6, 7). We and others have recently shown that TAK-981, a first-in-class inhibitor of the SUMO E1 activating enzyme, has potent anti-leukemic activity in AML preclinical models (8). In addition to a direct cytotoxic effect on the leukemic cells through the induction of their differentiation and death, inhibition of SUMOylation upregulates NK activating ligands (MICA/MICB and ICAM-1) at the surface of AML cells and favors their killing by NK cells (8). Along the same line, inhibition of the SUMO pathway upregulates Poliovirus Receptor (PVR), at the surface of multiple myeloma cells, favoring their recognition by DNAM1-activating receptor and killing by NK cells (9).

In addition to the overexpression of innate immunity activating ligands by cancer cells, recent studies have reported that inhibition of SUMOylation with TAK-981 activates an anti-tumor immune response and improves the efficiency of immune therapies by directly affecting immune cells. In models of pancreatic cancer and lymphomas, TAK-981 activates CD8 T-cells, NK cells, monocytes and dendritic cells (10–13), and decreases regulatory T-cells (T-reg) differentiation (14). The immuno-modulatory function of TAK-981 was suggested to rely on its ability to induce type-I Interferon (IFN-I) secretion. Blockade of IFN-I receptor (IFNAR) indeed blunts the anti-tumoral immune response induced by TAK-981 in different immunocompetent mouse cancer models (10, 12). Contrasting with these results, TAK-981-dependent activation of T cells from Chronic Lymphocytic Leukemia patients was found independent of IFNAR signaling (14).

Here, we demonstrate the potential of TAK-981 as a strategy to both restore the anti-leukemic activity of AML patients’ NK cells and increase the cytotoxicity of healthy donor’s NK and *ex vivo* expanded cord-blood NK. TAK-981 induces a strong increase in NK cells activation, degranulation capacity and inflammatory cytokines secretion. This enhances their cytotoxicity towards AML cell lines and patient blasts. At the molecular level, we demonstrate that the main effect of TAK-981 is to induce *IFNB1* expression and IFN-I secretion, which leads to autocrine activation of various ISGs and subsequent activation of NK cells. In addition, NK cells are activated *in trans* by IFN-I secreted by monocytes upon TAK-981 treatment.

## Results

### SUMOylation inhibition activates NK cells

To decipher whether inhibition of SUMOylation could activate NK cells, they were purified from PBMCs of healthy blood donors, treated with TAK-981 and co-cultured or not with either an AML cell line (U937, THP-1) or AML blasts purified from bone marrow of patients (Figure 1A). As expected, TAK-981 induced a fast deconjugation of both SUMO-1 and SUMO-2/3 from their targets after only 4 h (Figure 1B). TAK-981 had no effect on the viability of purified NK cells (Figure 1C, upper panel) or total Peripheral Blood Mononucleated Cells (PBMCs) (Figure 1C, lower panel) after 24 h of treatment. TAK-981 led to a strong overexpression of CD69, a marker of lymphocytes activation, on the surface of the purified NK cells co-cultured with U937 cells (Figure 1D). To confirm this observation *in vivo*, we then treated, SCID mice, which have no T- and B-lymphocytes but have normal NK cells, macrophages and granulocytes, with TAK-981 (Figure 1E). The activation of NK cells by TAK-981, as measured by the expression of CD69 (Figure 1F), was already significant after 5 h of treatment, maximal at 24 h and declined after 48 h (Figure 1G).

**Figure 1:**
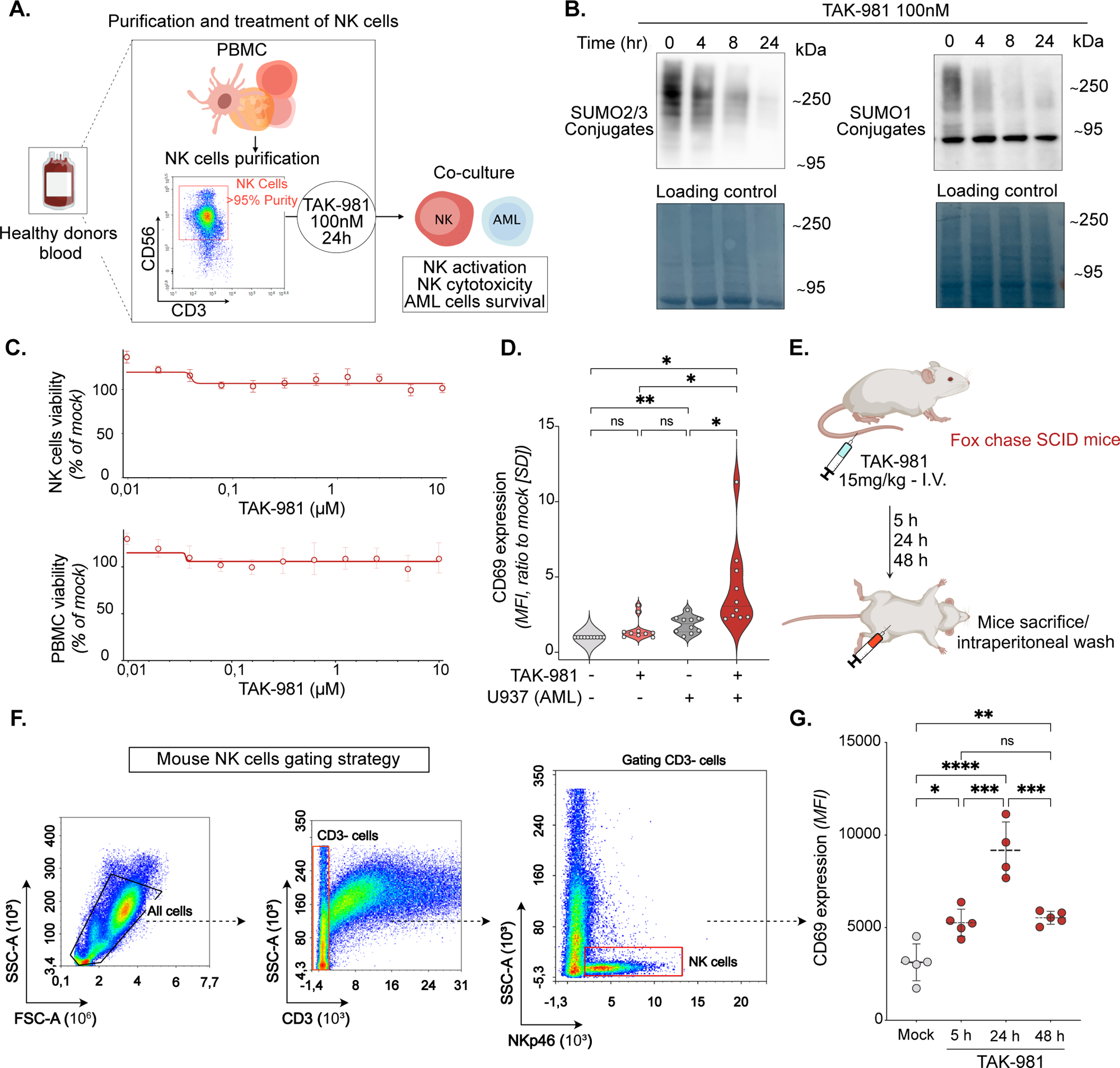
Inhibition of SUMOylation activates NK cells. **A.** Experimental procedure: primary NK cells were purified from healthy donors’ PBMCs. NK cells were treated with TAK-981 for 24 h and co-cultured or not with AML cells. When co-cultured, NK cells activation, cytotoxicity and AML survival were measured after 4 h of co-culture. **B.** Immunoblot with SUMO-1 and SUMO-2/3 antibodies of purified NK cells treated with 100 nM TAK-981 for the indicated times. Amido-black staining was used as loading control. **C.** NK cells (upper panel) or total PBMCs (lower panel) collected from (n=5) healthy donors were treated with increasing concentrations of TAK-981 for 24 h. Cell viability was determined by flow cytometry and a dose-response curve was generated by comparing the viability of TAK-981 treated cells with mock-treated controls. Data are shown as mean +/− SEM of 5 donors. **D.** Violin plot showing the expression of CD69 on the surface of purified NK cells. NK cells were treated with 100 nM TAK-981 for 24 h, followed by 4 h of co-culture with U937 (Effector:Target (E:T) ratio 1:1). Median fluorescent intensity of CD69 was normalized to mock-treated condition without co-culture (n= 10 donors, RM one-way ANOVA test). **E.** Fox chase SCID mice were injected intravenously with 15 mg/kg TAK-981 or vehicle. Mice were euthanized after 5h, 24h or 48h and intraperitoneal cavity wash was performed to collect NK cells. **F.** Gating strategies for mouse NK cells: total cells purified from the intraperitoneal cavity were stained with CD3 and NKp46 antibodies. NK cells were gated as SSC low CD3-NKp46+. G. CD69 expression levels on NK cells from SCID mice treated or not with TAK-981 for indicated time (4 or 5 mice per group, RM one-way ANOVA).

### SUMOylation inhibition increases NK cell cytotoxicity against AML

NK cell cytotoxicity requires formation of an immunological synapse with the target cells, leading to its degranulation and release of cytotoxic granules content. In the presence of U937 as target cells, the percentage of NK cells expressing the CD107a degranulation marker on their surface was higher when they were pre-treated with TAK-981 (Figure 2A and 2B). Confirming enhanced degranulation, TAK-981 stimulated the release of granzyme, perforin and granulysin from cytotoxic granules (Figure 2C). It also increased NK cells’ ability to secrete cytotoxic cytokines, among which TNFα and IFNγ (Figure 2C). Finally, TAK-981 induced the expression of *TNFSF10*, encoding for TRAIL, in NK cells (Figure 2D) and the subsequent increase of TRAIL at their surface (Figure 2E), which binds to its receptors (DR4 and DR5) on the surface of cancer cells to induce their apoptosis. Accordingly, TAK-981 increased the ability of NK cells to kill both U937 (Figure 2F, 2G and 2H) and THP-1 AML cell lines (Figure 2I and 2J) as well as AML patients blasts *in vitro* (Figure 2K).

**Figure 2:**
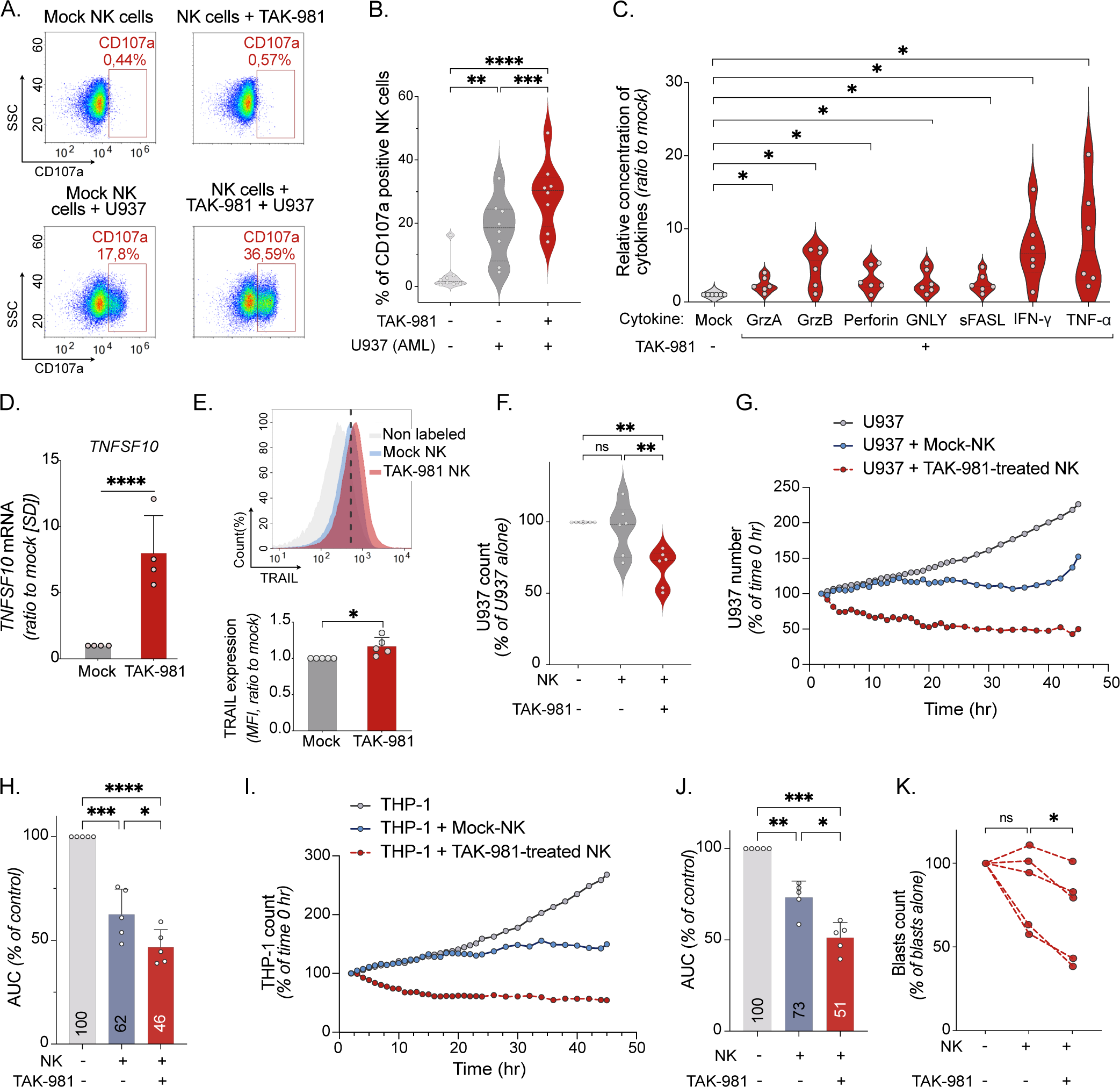
Inhibition of SUMOylation enhances NK cells cytotoxicity against AML. **A. B.** Mock- and TAK-981-treated NK cells were cocultured or not for 4 h with U937 cells. A. Flow cytometry profiles showing the percentage of CD107a positive NK cells. **B.** Quantification of the percentage of CD107a positive NK cells. (n=8 donors, RM one-way ANOVA test). **C.** LEGENDplex human NK panel was performed on supernatant of NK cells pre-treated or not with TAK-981 for 24 h and co-cultured for 4 h with U937 cells (E:T ratio 1:1). Relative quantity of each cytokine was normalized to mock-treated NK cells condition (n=6 donors, paired t-test). **D.** mRNA expression of *TNFSF10* in NK cells treated or not with TAK-981 for 24 h (n=4 donors, paired t-test). **E.** The expression of TRAIL on the surface of NK cells treated with TAK-981 for 24 h was measured by flow cytometry (upper panel) and presented as relative MFI (lower panel, n=5 donors, paired t-test). **F.** U937-LucZsGreen cells were co-cultured for 4 h with NK cells mock- or TAK-981-treated for 24 h. Relative number of U937 cells was measured by flow cytometry (LucZsGreen fluorescence) and normalized to U937 cells without NK cells co-culture (n= 6 donors, one-way ANOVA test). **G. H. I. J.** Real-time immune cell killing assay: purified NK cells were pre-treated or not with TAK-981 for 24 h, followed by co-culture with AML cell lines THP-1-LucZsGreen (G.) or U937-LucZsGreen (I.) at E:T ratio of 4:1. Green fluorescence was followed for 48 h using Incucyte device. **H.J.** THP-1 or U937 proliferation was calculated using the area under the curve for each co-culture condition (n= 5 donors, RM one-way ANOVA). **K.** AML patients’ blasts (5 patients) purified from bone marrow were co-cultured for 4 h with NK cells (n=3 donors) mock- or TAK-981-treated for 24 h (E:T ratio 1:1). Relative number of blast cells was measured by flow cytometry and normalized to blast cells number without NK cells co-culture.

### TAK-981 activates AML patients’ NK cells and *ex vivo* amplified NK, both *in vitro* and *in vivo*

To determine whether the inhibition of SUMOylation can also restore the activity of AML patient’s NK cells, we used PMBCs purified from the blood of AML patients at diagnosis. Treatment with TAK-981 increased their activation and degranulation capacity, as measured by the expression of CD69 (Figure 3A) and CD107a (Figure 3B), respectively. NK cells amplified from cord blood (eNK) are used for adoptive transfer in many clinical trials, including in AML (2, 15). Since eNK cells are expanded in the presence of IL-2 and IL-15, the levels of CD69 at their surface was higher than the basal level of expression on non-amplified NK cells from healthy donors (Figure 3C) and TAK-981 did not further increase its expression (Figure 3D). However, TAK-981 induced a progressive increase in their cytotoxicity, with a maximal activation after 48 h of treatment (Figure 3E). To decipher whether inhibiting SUMOylation could enhance the anti-leukemic activity of eNK cells *in vivo*, immunodeficient NSG mice grafted with THP-1 cells were injected with eNK cells and treated or not with TAK-981 (Figure 3F). Although both eNK cells and TAK-981 prolonged mice survival when used separately, we observed a much stronger effect on survival when mice were both injected with eNK cells and treated with TAK-981 (Figure 3G). Thus, targeting SUMOylation could be of therapeutic interest in AML to both restore the anti-leukemic activity of patient’s own NK cells and increase the cytotoxicity of allografted eNK cells.

**Figure 3:**
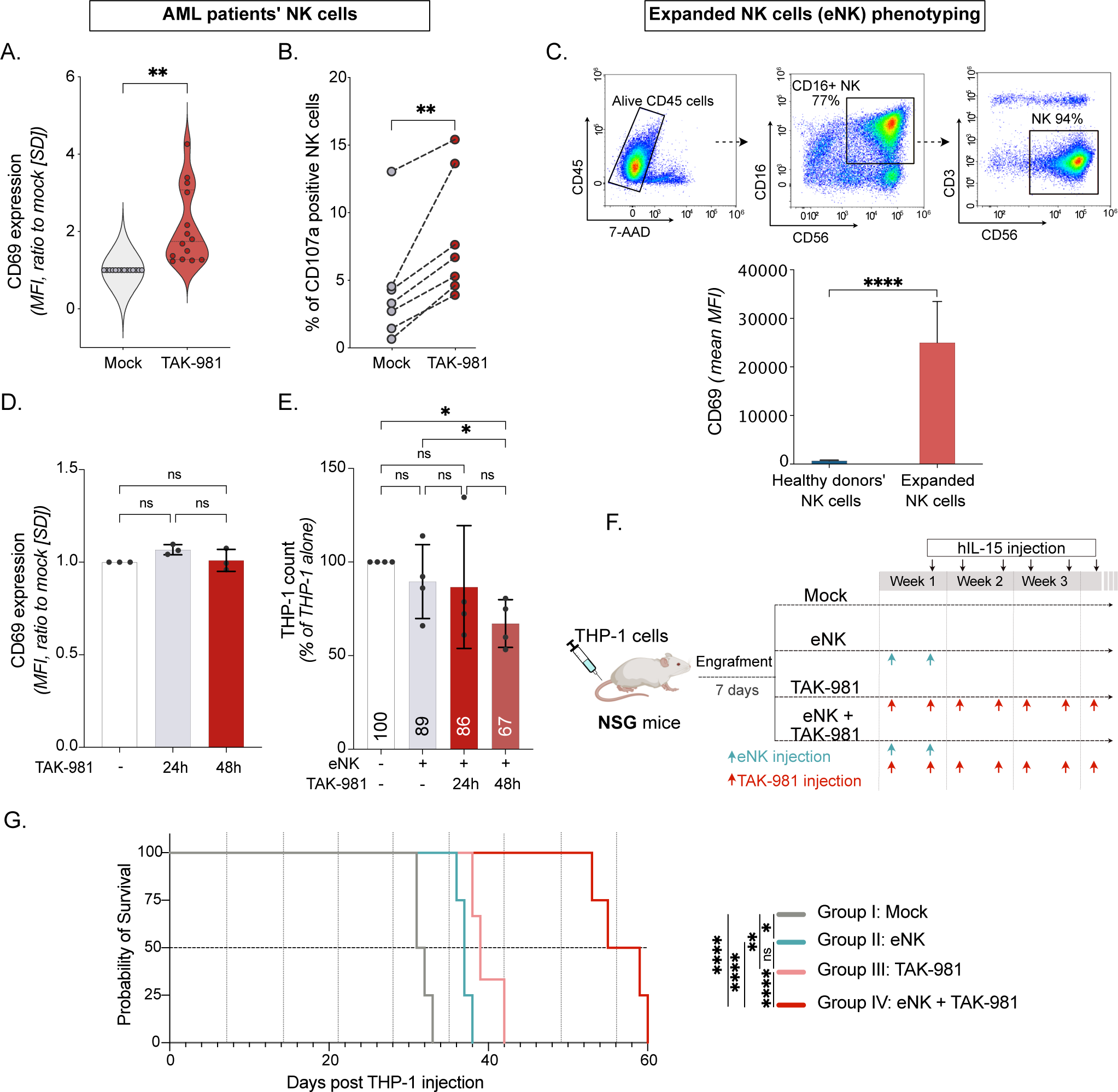
TAK-981 can both activate NK cells from AML patients and human eNK amplified from cord-blood. **A.** PMBCs purified from AML patient’s blood were treated with 100 nM TAK-981 for 24 h, and analyzed for CD69 expression. CD69 MFI on TAK-981-treated NK cells was normalized to mock-treated condition (n=14 patient samples, paired t test). **B.** Percentage of CD107a positive NK cells in AML patient’s PBMCs, treated or not with TAK-981 (n=7 patient samples, paired t-test). **C.** Upper panel, eNK cells phenotyping at day 14 of amplification. Lower panel, mean basal MFI of CD69 in NK cells purified from healthy donors compared to eNK (mean of n=11 for healthy donors’ NK, n=5 for eNK cells, unpaired t-test). **D.** eNK were treated with 100 nM TAK-981 for the indicated time and co-cultured with THP-1 cells for 4 h. CD69 MFI of TAK-981-treated eNK cells were normalized to mock-treated condition. Data are shown as mean +/− SD (n=3 eNK preparations, RM one way ANOVA). **E.** THP-1 cells were co-cultured for 4 h with mock- or TAK-981-treated eNK cells for the indicated times. The number of THP-1 cells was normalized to THP-1 cells without eNK cells co-culture. Data are shown as mean +/− SD (n=4 donors, RM one-way ANOVA). **F.** Experimental design: NSG mice were injected intravenously with THP-1-LucZsGreen cells. After THP-1 cells engraftment, mice were injected twice with eNK cells (day 7 and 10), followed by treatment with TAK-981 (15 mg/kg I.V.) and human recombinant IL-15 (hIL-15, 0.25 μg/mouse I.P.). For the rest of the experiment, mice were injected twice a week with TAK-981 and hIL-15. **G.** Overall survival was estimated in each group and compared with Kaplan-Meier method and log-rank test (n=4 for mock, n=3 for TAK-981, n=5 for eNK cells, n=4 for eNK+TAK-981).

### SUMOylation inhibition activates interferon type I response in NK cells

To decipher how SUMOylation controls NK cell activation, we performed RNA-Seq experiments on FACS-sorted NK cells from 3 different healthy donors treated with TAK-981 for 24 h. We identified 852 differentially expressed genes (log2 Fold change >1 or <-1, p<0.05), 586 being up-regulated and 266 down-regulated (Figure 4A, SupTable 1). Gene set enrichment analysis revealed a strong enrichment of pathways linked to inflammation and immune response, in particular the interferon response (Figure 4B and SupTable 1). Accordingly, TAK-981 induced the secretion of IFN-I by NK cells (Figure 4C). The secretion of IFN-I was necessary for the induction of Interferon Stimulated Genes (ISG) such as *OAS3* and *IFI44L,* as it was prevented by blocking the IFN-I receptor IFNAR with the MMHAR2 antibody (Figure 4D). *IFNB1* induction was not affected by the IFNAR blocking (Figure 4E), suggesting its direct regulation by SUMOylation.

**Figure 4:**
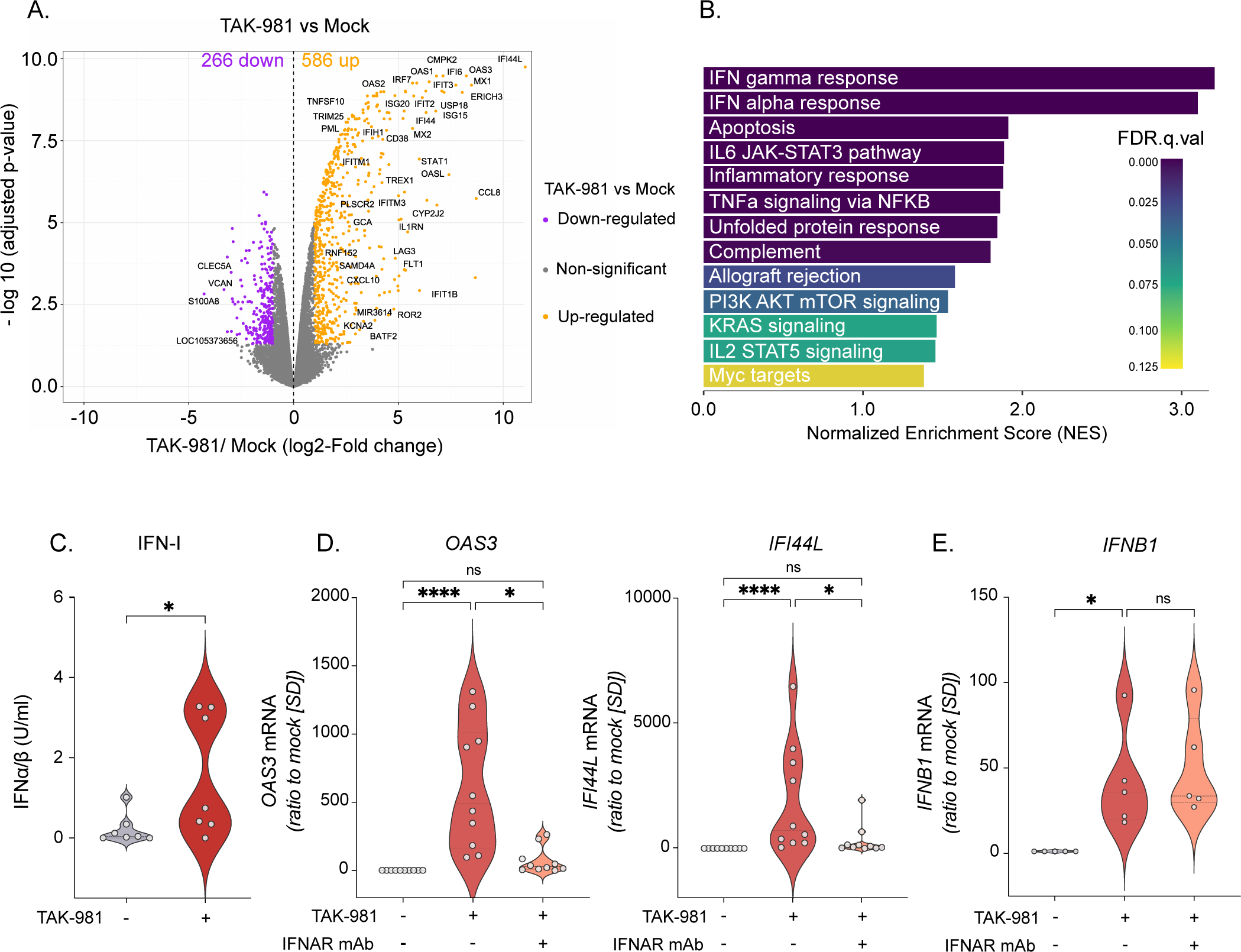
Targeting SUMOylation induces interferon pathways in NK cells. **A.** Volcano plot of differentially expressed genes analyzed by RNA-Seq in NK cells purified from healthy donors’ blood (n=3) treated for 24 h with 100 nM TAK-981 compared to mock-treated controls. **B.** Gene Set Enrichment Analysis (GSEA) on Hallmark datasets on the RNA-Seq data presented in A. All pathways significantly enriched in TAK-981-compared to mock-treated cells are shown (abs(NES) > 1, p < 0.05 and FDR < 0.25). **C.** Relative concentration of IFN-I in the supernatant of NK cells purified from healthy donors’ blood (n=7 donors) and treated with TAK-981 for 24 h. IFN-I was quantified with HEK blue IFN-α/β reporter cells (paired t-test). **D. E.** mRNA expression levels of *OAS-3*, *IFI44L* (D) and *IFNB1* (E) in NK cells treated or not with TAK-981 and IFN-I receptor (IFNAR) neutralizing monoclonal antibody for 24 h (n=10 donors for *OAS-3* and *IFI44L*, n=5 donors for *IFNB1.* RM one-way ANOVA).

### SUMOylation inhibition activates NK cells through the induction of *IFNB1*

The secretion of IFN-I was required for TAK-981-induced activation of NK cells, as blocking IFNAR prevented TAK-981-induced expression of CD69 at their surface (Figure 5A). Treatment of C57BL/6 mice with TAK-981 induced the activation of NK cells in their blood (Figure 5B and 5C) and spleen (Figure 5D) after 24 h. This was not the case for NK cells from spleen and blood of mice knock-out for IFNAR (Figure 5D). Although TAK-981 did not activate NK cells in the bone marrow of wild-type mice (Figure 5E), it increased their numbers in this compartment in wild-type but not in IFNAR knock-out mice (Figure 5F). Altogether, this suggests that one main effect of TAK-981 in NK cells is to induce *IFNB1*. The subsequent secretion of IFN-I increases the transcription of various ISGs, which contributes to the activation of NK cells both *in vitro* and *in vivo*.

**Figure 5:**
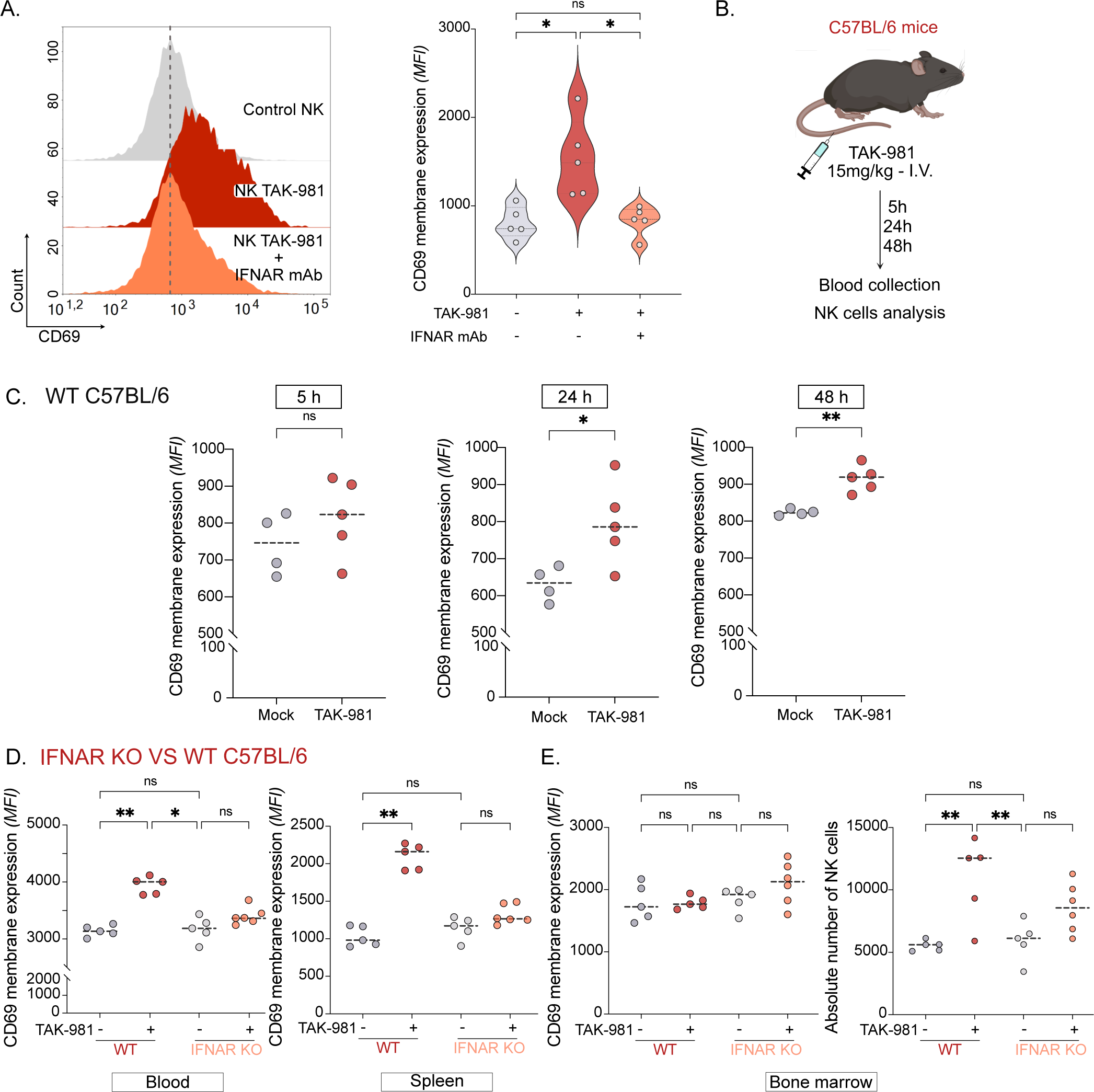
TAK-981-induced secretion of IFN-I is required for NK cells activation. **A.** MFI of CD69 expression on the surface of NK cells treated or not with TAK-981 and IFNAR neutralizing monoclonal antibody for 24 h (left panel). Violin plot showing the quantification of CD69 expression (MFI) on the surface of NK cells treated or not with TAK-981 and IFNAR neutralizing monoclonal antibody for 24 h (right panel, n=5 donors, RM one-way ANOVA). **B.** C57BL/6 mice were injected intravenously with 15 mg/kg TAK-981 or vehicle. Blood was collected after 5 h, 24 h and 48 h and NK cells activation was analyzed. **C.** CD69 expression levels (MFI) on NK cells from blood of mice treated or not with TAK-981 (4 or 5 mice per group, RM one-way ANOVA). **D.** WT and IFNAR KO C57BL/6 mice were injected intravenously with 15 mg/kg TAK-981 or vehicle. Mice were euthanized after 24 h and NK cells from blood, spleen and bone marrow were analyzed. CD69 expression levels on NK cells from blood and spleen of WT and IFNAR KO mice treated or not with TAK-981 (5 or 6 mice per group, RM one-way ANOVA). **E.** CD69 expression levels on NK cells and their absolute number per mice in bone marrow of WT and IFNAR KO mice treated or not with TAK-981 (5 or 6 mice per group, RM one-way ANOVA).

### NK cells are also activated *in trans* by IFN-I secreted by myeloid cells upon inhibition of SUMOylation

NK cells present within PBMCs were more activated by TAK-981 than purified NK cells (Figure 6A). Within the PBMCs, NK cells were however the main cells responsible for TAK-981-induced cytotoxicity on U937 cells as (i) NK cells purified from TAK-981 treated PBMCs had equivalent cytotoxicity as total PBMCs (ii) and their depletion from PBMCs prevented U937 lysis (Figure 6B). Transwell experiments showed that soluble factors secreted by TAK-981-treated PBMCs could indeed activate NK cells and increase their cytotoxicity. Moreover, blocking IFNAR prevented their activation by these soluble factors, suggesting a main role for IFN-I in NK cells activation (Figure 6C, D). To identify, which immune cells, in addition to NK cells, are producing IFN-I upon inhibition of SUMOylation, T lymphocytes, B lymphocytes, NK cells, monocytes and dendritic cells were FACS-sorted from the PBMCs of 5 different healthy blood donors. Purified cells were then treated *in vitro* with TAK-981 and IFN-I secretion was monitored using a reporter assay. This revealed that monocytes are by far the strongest producers of IFN-I upon TAK-981 treatment (Figure 6E). *IFNB1* was induced by TAK-981 to a mean of 1800-fold in purified monocytes and to a mean of 35-fold in NK cells (Figure 6F). TAK-981 also strongly induced ISGs such as *OAS3* and *IFI44L* in monocytes (Figure 6G). Finally, although maximal at 24 h of treatment, the expression of *IFNB1* in monocytes was already highly increased after 4 h and, similar to NK, was not affected by IFNAR blocking (Figure 6H). Altogether, this suggests that inhibition of SUMOylation can also activate NK cells *in trans,* through the induction of IFN-I secretion by monocytes.

**Figure 6:**
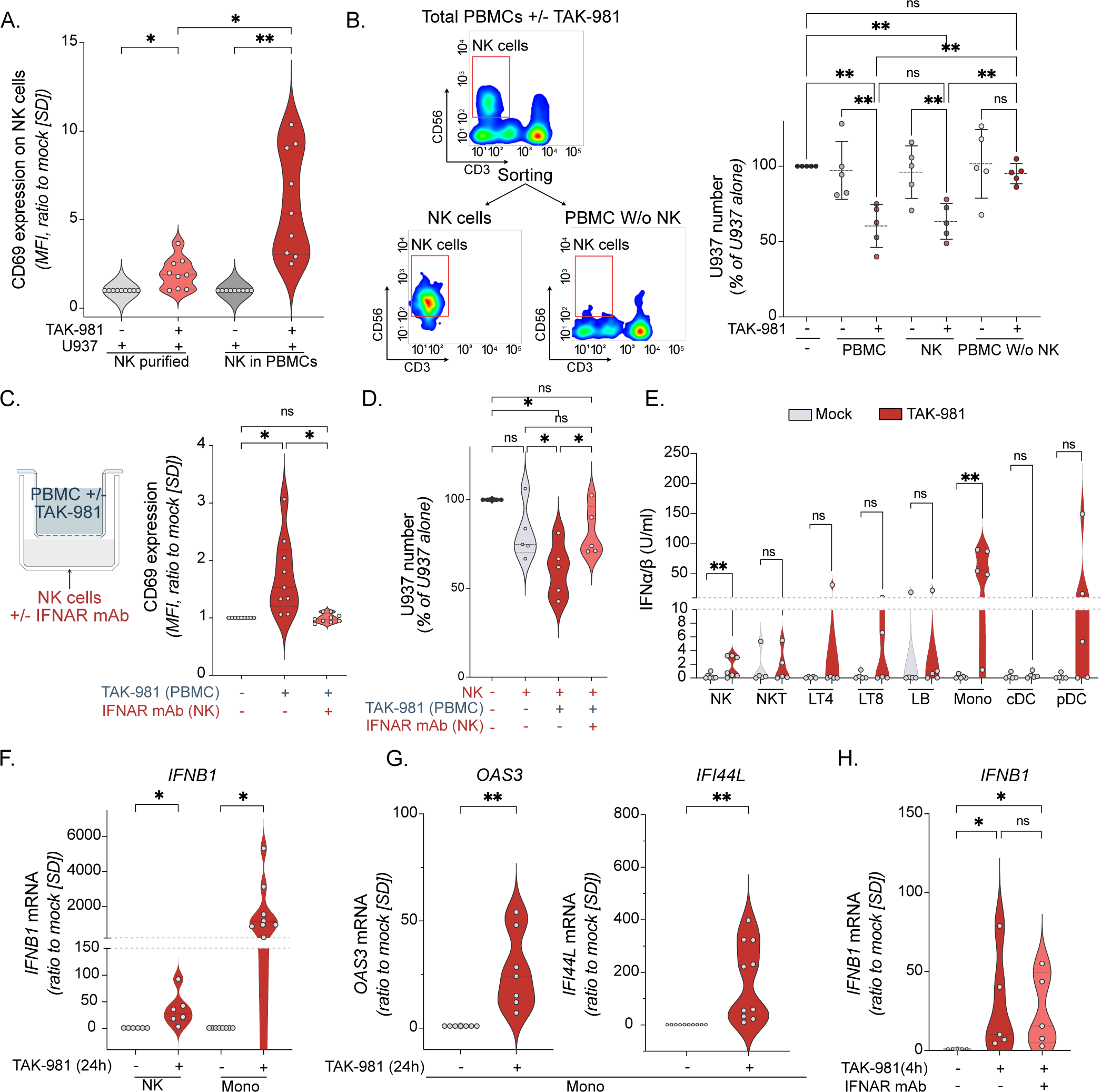
NK cells are activated *in trans* by IFN-I secreted by myeloid cells upon inhibition of SUMOylation. **A.** Purified NK cells or PBMCs were treated with 100 nM TAK-981 for 24 h, followed by 4 h of co-culture with U937 (E:T ratio 1:1). CD69 expression (MFI) on NK cells was normalized to mock-treated condition (n=8 donors, RM one-way ANOVA). **B.** Total PBMCs purified from healthy donors’ blood were treated with TAK-981 for 24 h followed by FACS to separate NK cells (CD56+ CD3) and NK-depleted PBMCs. Sorted populations or total PBMCs where co-cultured for 4 h with U937-LucZsGreen cells. Relative number of U937 cells was measured by flow cytometry (LucZsGreen expression) and represented as the percentage of U937 cells without NK cells co-culture (n=5 donors, RM one-way ANOVA). **C.** PBMCs were treated with TAK-981 for 24 h, washed and transferred in the upper chamber of a transwell. Purified NK cells +/− IFNAR blocking monoclonal antibody were placed in the lower chamber. After 24 h, NK cells were co-cultured with U937-LucZsGreen cells for 4 h. CD69 expression was measured on the surface of NK cells and normalized to mock-treated condition (right panel, n=9 donors, RM one-way ANOVA test). **D.** Relative number of U937 cells after 4 h of co-culture with NK cells treated as in C (n=5 donors, RM one-way ANOVA). **E.** NK cells, NKT cells, CD4+T cells (LT4), CD8+T cells (LT8), B-lymphocytes (LB), monocytes (Mono), classical dendritic cells (cDC) and plasmacytoid dendritic cells (pDC) were FACS-sorted and treated with TAK-981 for 24 h. IFN-I concentration was quantified in the supernatant with HEK Blue IFN-α/β reporter cells (n=5 donors, paired t-test). **F.** Relative expression of IFNB1 mRNA in NK cells vs monocytes purified from healthy donors’ blood and treated for 24 h with TAK-981. (n=6 donors for NK cells, and n=8 donors for monocytes, paired t-test). **G.** Relative expression of OAS-3 and IFI44L mRNA in purified monocytes after 24 h treatment with TAK-981 (n=7 donors for OAS-3 gene, n=10 donors for IFI44L gene, paired t-test). **H.** Relative expression levels of IFNB1 mRNA in purified monocytes treated for 4 h with TAK-981 +/− IFNAR blocking monoclonal antibody (n=5 donors, paired t-test).

## Discussion

NK cells are increasingly considered as attractive therapeutic tools for cancer therapies either through the activation of patient’s own NK cells or by adoptive transfer of *in vitro* expanded NK cells (16). Here, we show that NK cell activation is under the control of the SUMO pathway and targeting SUMOylation with TAK-981 increases their anti-tumoral activity. We found that one main effect of SUMOylation in NK cells is to repress IFN-I pathway. Inhibition of SUMOylation with TAK-981 indeed up-regulated more than 500 genes after 24 h, most of them belonging to the interferon pathway. The strong TAK-981-induced up-regulation of ISGs was abolished when the IFN-I receptor (IFNAR) was blocked, suggesting that the secretion of IFN-I is required for their expression. The induction of the *IFNB1* gene by TAK-981 was not dependent on prior IFN-I secretion, suggesting it constitutes the primary target of TAK-981 in NK cells. The spontaneous induction of *IFNB1* was observed in various cell types upon inhibition of SUMOylation, either by knocking-out the SUMO E1 or E2 enzymes (17, 18) or upon TAK-981 treatment (10, 11, 19). How SUMOylation controls *IFNB1* is however still largely unknown. Interferon response factors IRF-1, IRF-2, IRF-3 and IRF-7 are SUMOylated and their SUMOylation limits their activity as measured by luciferase reporter assays (20–24). The inhibition of their SUMOylation by mutation of their SUMOylated lysines increased the induction of *IFNB1* following viral infection but had however no effect on the basal *IFNB1* expression (23), suggesting that deSUMOylation of IRFs is not sufficient to induce *IFNB1*. Other upstream regulators of *IFNB1* are SUMOylated. This is the case of MDA5, MAVS and RIG-I, whose SUMOylation was suggested to rather increase *IFNB1* expression (25–28). Altogether, this suggests that TAK-981 induced expression of *IFNB1* does not rely on the deSUMOylation of the above-mentioned proteins. The proteins, whose deSUMOylation is sufficient to induce *IFNB1* could be those bound to the cis-regulatory regions controlling its expression. Indeed, in mouse bone marrow derived dendritic cells (BMDCs), yet non-identified SUMOylated proteins were found enriched on 3 enhancers located between 15 and 30 kb from the gene body and suggested to participate in *IFNB1* repression (18). However, these enhancers are not conserved in human cells. Hence, the molecular mechanisms underlying the strong and rapid induction of *IFNB* by TAK-981 remain to be identified.

TAK-981-induced IFN-I response was suggested to play a critical role in its anti-tumoral activity through its ability to activate both innate and adaptative anti-tumoral immune responses (10–13, 29). Accordingly, TAK-981 was found to enhance the antitumoral activity of monoclonal antibodies as seen for rituximab (anti-CD20) or daratumumab (anti CD38) in preclinical mouse models of lymphoma and myeloma respectively (11). These combination therapies are being evaluated in two clinical trials (NCT04776018 and NCT04074330). Here, we show that TAK-981 activates the cytotoxicity of human NK cells towards AML cells both *in vitro* and *in vivo*. As mentioned, this relies on IFN-I secreted by the NK cells themselves, but also by monocytes, which were by far the strongest TAK-981-induced IFN-I producers in the blood. As a consequence of IFN-I signaling activation, TAK-981 largely increased the degranulation capacity of NK cells as well as their ability to secrete cytotoxic cytokines such as TNF-α, IFN-γ and TRAIL. AML patients are generally treated with an immunosuppressive chemotherapy, which leads to bone marrow aplasia characterized by a loss of immune cells, including NK cells. At complete remission (CR), NK cell number increases but not to levels present in healthy donors. Moreover, although their degranulation capacities are restored at CR, their ability to produce cytokines remains low (30). As we demonstrate that TAK-981 can activate AML patient’s NK cells, it could be used after CR to increase NK cells activity towards residual AML cells. TAK-981 would thus have a dual anti-leukemic effect by targeting AML cells directly as we and others have recently shown (8, 31) and indirectly *via* the activation of patient’s own NK cells. Altogether, this would maximize the chances to control minimal residual disease (MRD), which is largely responsible for the high relapse rate in AML. Finally, we report that TAK-981 increases the cytotoxicity of *ex vivo* expanded NK cells from haplo-identical cord blood donors (eNK), both *in vitro* and *in vivo.* In contrast to NK cells isolated from adult donors PBMCs, NK cells from cord blood can be efficiently expanded and matured *ex vivo*, making them ideal off-the-shelf cellular therapy (16, 32). Due to their anti-tumoral efficacy and their low toxicity, adoptive transfer of eNK cells is increasingly considered for therapeutic use, including in AML (15, 16). Hence, TAK-981 could be used to further increase the anti-tumoral efficacy of eNK cell-based therapies.

## Methods

### AML patients and healthy donor samples

Buffy coats were obtained from healthy blood donors from the Etablissement Français du Sang (EFS, agreement n°21PLER2019-0002). For AML patient samples, bone marrow aspirates and blood samples were collected after obtaining written informed consent from patients under the frame of the Declaration of Helsinki and after approval by the Institutional Review Board (Ethical Committee “Sud Méditerranée 1,” ref 2013-A00260-45, HEMODIAG_2020 cohort).

### Myeloid and NK cells purification

PBMCs were purified from either buffy coat of healthy donors or blood and bone marrow of AML patients using density-based centrifugation (Histopaque 1077-H8889 Sigma-Aldrich).

Myeloid cells were purified from PMBCs using CD33+ microbeads (Miltenyi Biotech 130-045-501) according to manufacturer guidelines. The flow through, depleted from CD33+ cells, was then processed using negative selection NK cells isolation Kit (Miltenyi Biotech 130-092-657). NK cells purity was assessed by flow cytometry using CD56 and CD3 expression markers.

### FACS sorting of immune cells populations

PBMCs were washed once with PBS, incubated 15 min at room temperature in PBS + 2% Fetal Bovine Serum (FBS) with conjugated antibodies (see table 1); washed with PBS and sorted using a BD FACS Aria IIu Cell sorter. Cell populations were collected in PBS + 2% FBS before returning to culture medium (RPMI + 10% FBS). The following markers were used for sorting the different populations: NK cells: CD56+/CD3−; NKT cells: CD56+/CD3+; CD8 T lymphocytes: CD3+/CD8+; CD4 T lymphocytes: CD3+/CD4+; B lymphocytes: CD3−/CD19+; Monocytes HLA-DR+/CD14+; Conventional dendritic cells (cDC): HLA-DR+/CD14−/CD11c+; Plasmacytoid dendritic cells (pDC): HLA-DR+/CD14−/CD11C−/CD123+. For NK cells depletion experiments, NK cells (CD56+/CD3-) were separated from the rest of PBMCs using BD FACS Aria IIu Cell sorter.

**Table 1:**
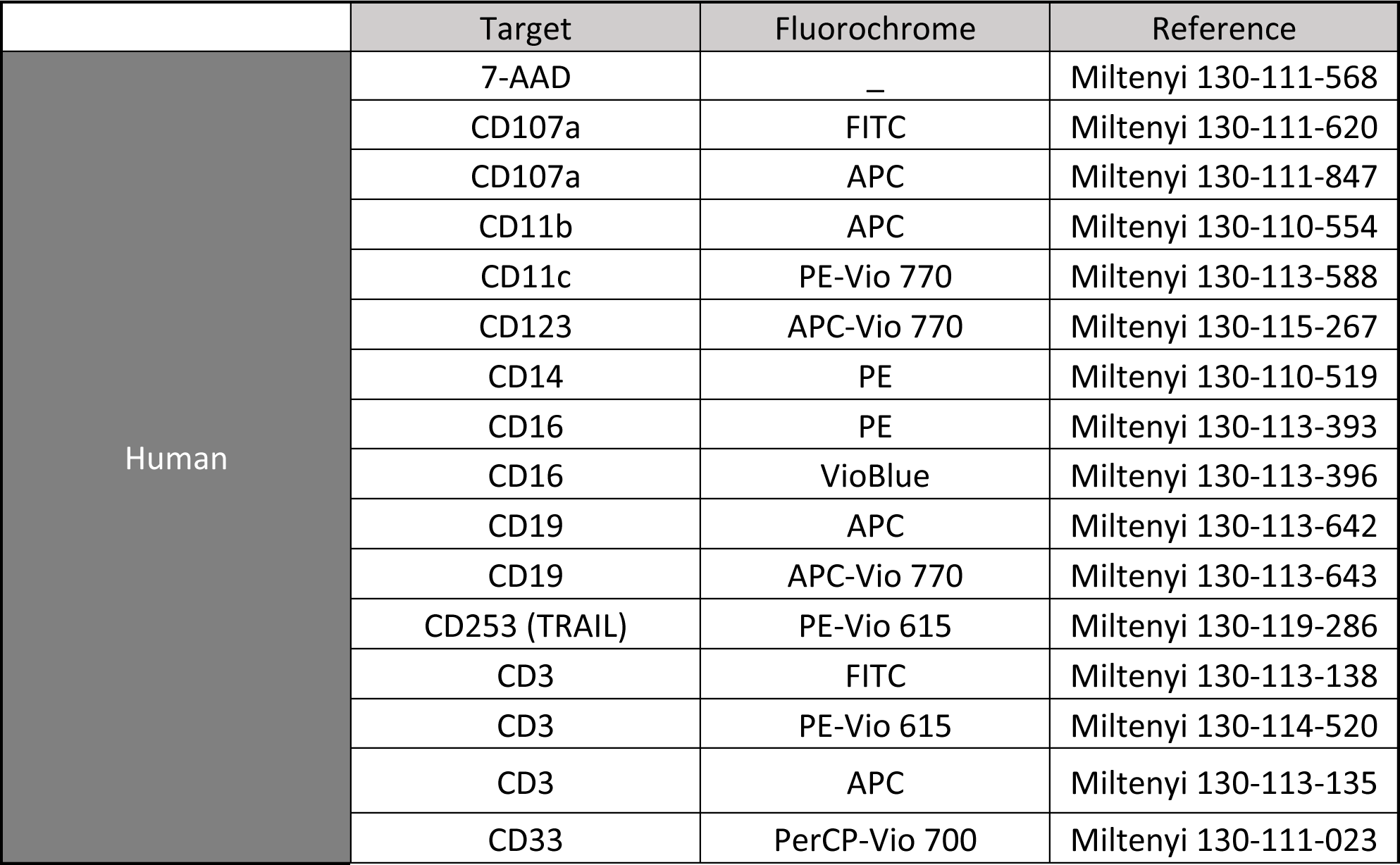
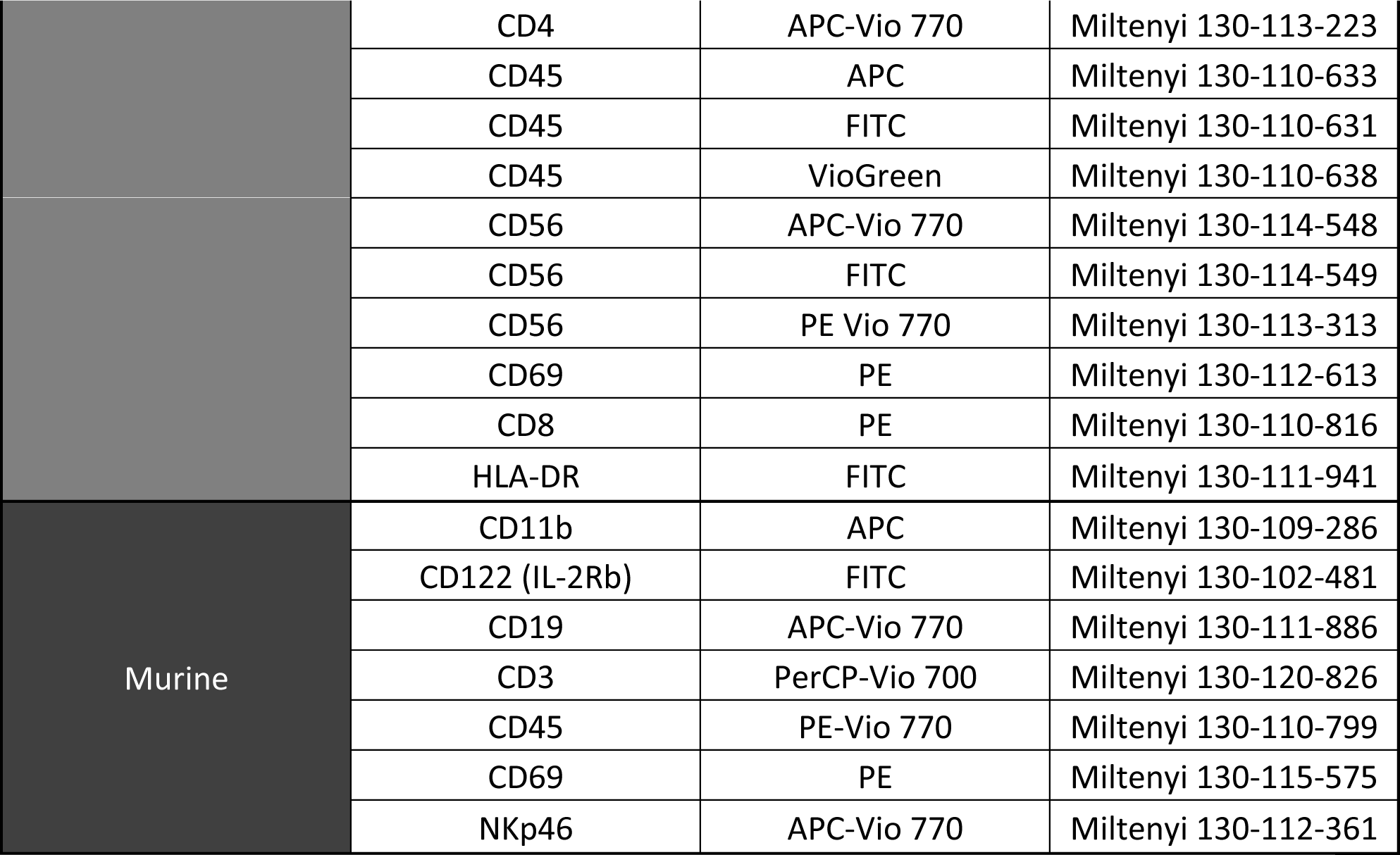
List of antibodies used for flow cytometry.

|  | Target | Fluorochrome | Reference |
| --- | --- | --- | --- |
| Human | 7-AAD | — | Miltenyi 130-111-568 |
|  | CD107a | FITC | Miltenyi 130-111-620 |
|  | CD107a | APC | Miltenyi 130-111-847 |
|  | CD11b | APC | Miltenyi 130-110-554 |
|  | CD11c | PE-Vio 770 | Miltenyi 130-113-588 |
|  | CD123 | APC-Vio 770 | Miltenyi 130-115-267 |
|  | CD14 | PE | Miltenyi 130-110-519 |
|  | CD16 | PE | Miltenyi 130-113-393 |
|  | CD16 | VioBlue | Miltenyi 130-113-396 |
|  | CD19 | APC | Miltenyi 130-113-642 |
|  | CD19 | APC-Vio 770 | Miltenyi 130-113-643 |
|  | CD253 (TRAIL) | PE-Vio 615 | Miltenyi 130-119-286 |
|  | CD3 | FITC | Miltenyi 130-113-138 |
|  | CD3 | PE-Vio 615 | Miltenyi 130-114-520 |
|  | CD3 | APC | Miltenyi 130-113-135 |
|  | CD33 | PerCP-Vio 700 | Miltenyi 130-111-023 |
|  | CD4 | APC-Vio 770 | Miltenyi 130-113-223 |
|  | CD45 | APC | Miltenyi 130-110-633 |
|  | CD45 | FITC | Miltenyi 130-110-631 |
|  | CD45 | VioGreen | Miltenyi 130-110-638 |
|  | CD56 | APC-Vio 770 | Miltenyi 130-114-548 |
|  | CD56 | FITC | Miltenyi 130-114-549 |
|  | CD56 | PE Vio 770 | Miltenyi 130-113-313 |
|  | CD69 | PE | Miltenyi 130-112-613 |
|  | CD8 | PE | Miltenyi 130-110-816 |
|  | HLA-DR | FITC | Miltenyi 130-111-941 |
| Murine | CD11b | APC | Miltenyi 130-109-286 |
|  | CD122 (IL-2Rb) | FITC | Miltenyi 130-102-481 |
|  | CD19 | APC-Vio 770 | Miltenyi 130-111-886 |
|  | CD3 | PerCP-Vio 700 | Miltenyi 130-120-826 |
|  | CD45 | PE-Vio 770 | Miltenyi 130-110-799 |
|  | CD69 | PE | Miltenyi 130-115-575 |
|  | NKp46 | APC-Vio 770 | Miltenyi 130-112-361 |

### Cell culture

Human AML cell lines U937-LucZsGreen and THP-1-LucZsGreen express both luciferase and ZsGreen protein (8). AML cell lines and human primary cells were cultured with RPMI 1640 (R8758 Sigma-Aldrich) medium complemented with 10% FBS, Penicillin and Streptomycin, at 37°C / 5% CO2. IFN-α/β Reporter HEK 293 Cells were obtained from InvivoGen and cultured in DMEM medium (Gibco-41965-039) complemented with 10% FBS, Penicillin, Streptomycin, Blasticidin and Zeocin at 37°C / 5% CO2.

### eNK amplification

eNK expansion protocol was adapted from a previously described protocol (33) using umbilical cord blood (UCB) (biobank BB-0033-00031 from CHU Montpellier). Briefly, CD3 positive cells were removed from UCBMCs using EasySep™ Human CD3 Positive Selection Kit II (Stemcell), according to manufacturer’s instructions. CD3 negative cells were cultured at 1×10^6^ cells/mL in eNK medium (RPMI1640 Glutamax™ or NK MACS, Miltenyi) supplemented with 10% FBS or 5% human serum, 100 IU/mL rhIL-2 (Peprotech) and 5 ng/mL hrIL-15 (Miltenyi). From day 5-7 and every 2-3 days until day 14-21, medium was removed and freshly prepared eNK medium added to reach 0.6×10^6^ cells/mL. 70 Gy-irradiated lymphoblastoid B-EBV feeder cells at appropriate ratios were added.

At the end of expansion, eNK cells phenotype was analyzed by flow cytometry at day 14. 2.10^5^ cells were stained in PBS + 2% FBS with 7AAD (Viability), anti-CD45-VioGreen, anti-CD19-APC-Vio770, anti-CD56-PE-Vio770, anti-CD69-PE, anti-CD16-VioBlue, anti-CD3-APC. Cells were incubated for 20 min at 4°C in the dark, followed by 2 washes using PBS containing 2% FBS. Cells were then resuspended in PBS + 2% FBS and acquired on a Gallios flow cytometer (Beckman Coulter).

### Pharmacological inhibitors and survival analysis

TAK-981 was obtained from Takeda Development Center Americas, Inc. TAK-981 was re-suspended in DMSO at 10 mM and used at 100 nM. IFNAR2 neutralizing monoclonal antibody (MMHAR2 Sigma-Aldrich 407295) was used at 1 µg/ml.

To measure the IC_50_ of TAK-981, total PBMCs were treated with increasing concentrations [10 nM -> 10 µM] of TAK-981 for 24 h. Cell survival was measured using flow cytometry based on the FSC and SSC parameters for total PBMCs, and using CD56 and CD3 markers to identify NK cells.

### NK cells cytotoxicity assay

Purified NK cells or total PBMCs from healthy donors were treated with TAK-981 or mock for 24 h. After treatment, cell media was changed and cells were co-cultured with AML target cells (AML patient’s blasts, U937-LucZsGreen or THP-1-LucZsGreen at a ratio of effector:target cells of 1:1 for NK cells or 10:1 for PBMCs), unless otherwise indicated. AML patient’s blasts previously purified from bone marrow were thawed and stained with 1 µM CFSE (eBioscience-65-0850-85) prior to co-culture. Four hours after co-culture, NK cells activity was measured by flow cytometry using CD69 and CD107a markers and the total number of green fluorescent AML cells was counted.

To follow the cytotoxic activity of NK cells against AML cell lines in real time, the number of fluorescent AML cells was assessed during 48 h using the IncuCyte S3 Live Cell imaging system (Sartorius) in the non-adherent cell-by-cell mode 20X objective, 9 images/well and 1 image/hour. Analyses were performed by measuring the relative integrated green fluorescence intensity using IncuCyte 2022A software.

### Flow cytometry

For NK cells degranulation assay, CD107a antibody was added to the cells at the beginning of the co-culture for healthy donors or 4 h before staining for AML patients’ PBMCs. One hour later, Monensin sodium salt (Sigma-Aldrich M5273) was added at 6 µg/ml to block endocytosis. Cells were then incubated for 3 h before being analyzed with NovoCyte Flow Cytometer (Agilent). For the other markers, cells were washed once with PBS, incubated 15 min at room temperature in PBS 5% FBS with conjugated antibodies (see table 1), washed with PBS and analyzed with NovoCyte Flow Cytometer (Agilent). Median fluorescence intensities (MFI) or the percentages of positive cells were calculated using the NovoExpress software (v.1.5.6).

### Cytokines quantification

Mock or TAK-981-treated NK cells purified from healthy donors were co-cultured for 4 h with U937 cells at a ratio of effector:target cell of 1:1. Supernatants were collected and cytokines were quantified using Human CD8/NK panel (LEGENDplex 740267) according to manufacturer guidelines.

IFN-I was quantified using IFN-α/β reporter HEK 293 cells. Cells were seeded at 30.000 cells/well of 96-well plate. After 24 h, cell culture supernatants were added to the reporter cells and incubated overnight. SEAP activity was then quantified using Quanti-Blue solution (InvivoGen, rep-qbs) and absorbance was measured using FLUOstar Omega device (BMG Labtech).

### Transwell assay

PBMCs were treated for 24 h with mock or 100 nM TAK-981. After treatment PBMCs were placed in the upper chamber of transwell inserts with 0.4 µm pore size (Falcon - 353095) and NK cells, treated or not with MMHAR2, were placed in the lower chamber. NK cells were collected 24 h later and co-cultured with U937-LucZsGreen cells for 4 h. CD69 expression on NK cells and AML cells survival were analyzed by flow cytometry.

### *In-vivo* mice models

All experiments on animals were approved by the Ethics Committee of the Languedoc-Roussillon (2018043021198029 #14905 v3). As specified by the Ethical Committee, mice were observed daily for signs of distress (ruffled coat, hunched back, and reduced mobility). These observations were used to determine the appropriate time for euthanizing the animals.

6-10 weeks old Fox chase mice (CB17/Icr-Prkdc^scid^/IcrIcoCrl) were treated IV with TAK-981 (15 mg/kg) resuspended in 20% HPBCD (hydroxypropyl beta-cyclodextrin) or vehicle (20% HPBCD). Mice were euthanized after 5 h, 24 h or 48 h of treatment. Intraperitoneal wash was performed by injecting 5 mL of PBS + 3% FBS into the peritoneal cavity of the mice followed by recollecting the PBS. Total cells collected were labeled with CD3, CD69 and NKp46 antibodies (table 1) and analyzed by flow cytometry.

To determine the kinetics of activation of NK cells by TAK-981 *in vivo*, C57BL/6J female mice (6-10 weeks old) were treated IV with TAK-981 (15 mg/kg) or vehicle. Blood was collected after 5 h, 24 h or 48 h of treatment. In the same manner, C57BL/6J WT or IFNRA KO mice were treated or not for 24 h with TAK-981 then euthanized. Blood, spleen and bone marrow (from femurs) were collected and cells were dissociated and washed with PBS. Red blood cells were lysed using the ACK lysis buffer (A1049201, Gibco) and mononuclear cells were labeled with CD11b, CD19, CD69, CD3, CD45, CD122 (table 1) and analyzed by flow cytometry. Mice NK cells were gated as CD45+/CD11b-SSC low/CD3−/CD19−/CD122+, and NK cells activation level was measured using Median fluorescent Intensity (MFI) of CD69.

For Cell Line Derived Xenograft (CLDX), 6-10 weeks old female NOD/LtSz-SCID/IL-2Rγchain null (NSG) mice (Charles River) were treated IV with 30 mg/kg busulfan (SIGMA B2635). Two days after, mice were injected IV with 10^6^ THP-1-LucZsGreen cells resuspended in PBS. Mice were then injected with 10^7^ expanded NK cells (eNK) at day 7 and 10 post AML injection followed by another injection at day 10. Mice were then treated with TAK-981 (IV, 15 mg/kg) and IL-15 (IP, 0.25 µg/mice, Miltenyi Biotech 130-095-760) twice a week.

### RNA-seq libraries preparation and sequencing

FACS-sorted NK cells sorted from 3 different healthy donors’ blood were treated with mock or 100 nM TAK-981 for 24 h. Total RNA was purified using GenElute Single Cell RNA Purification Kit (Sigma-Aldrich RNB300), treated with DNase I on column (Sigma-DNASE70-1SET). RNA quality was checked using a BioAnalyser Nano 6000 Chip (Agilent). Libraries were prepared using SMART-Seq Stranded Kit (Takara Bio). The quality, size and concentration of cDNA libraries were checked using the High Sensitivity NGS kit Fragment Analyzer and qPCR (ROCHE Light Cycler 480). Libraries were sequenced using an Illumina Novaseq 6000 sequencer as single-end 100 base reads. Image analysis and base calling were performed using the NovaSeq Control Software, Real-Time Analysis 3 (RTA) and bcl2fastq. The RNA-Seq sequencing data are available on Gene Expression Omnibus with accession number GSE255279.

### RNA-seq mapping, quantification and differential analysis

RNA-seq reads were mapped on the Human reference genome (hg38, GRCh38p12) using TopHat2 (2.1.1)(34) based on the Bowtie2 (2.3.5.1) aligner (35). Reads association with annotated gene regions was done using the HTseq-count tool v0.11.1 (36). Differential expression analysis was performed with edgeR using counts per million normalization and quasi-likelihood F-test with paired design (37). Genes with a fold change ≥2 or ≤ 0.5 and an adjusted p-value < 0.05 were considered differentially expressed. Gene Set Enrichment Analyses were performed using https://www.gsea-msigdb.org/gsea/index.jsp (version 4.0.3)(38).

### RT-qPCR assays

Total RNA was purified using TRIzol reagent (Invitrogen - 15596026), after DNase I treatment with RQ1 RNase-Free DNase (Promega - M6101), reverse transcription was done using OneScript Plus cDNA Synthesis Kit (abm-G236). qPCR was then assayed on 0.2 µg of cDNA using Platinum Taq DNA polymerase (Invitrogen-3644073) and specific DNA primers (IDT sequences in table 2). qPCR reaction was analyzed on CFX opus Real-Time PCR device (Bio-Rad) and data were normalized to the expression levels of *GAPDH* as housekeeping gene.

**Table 2:**
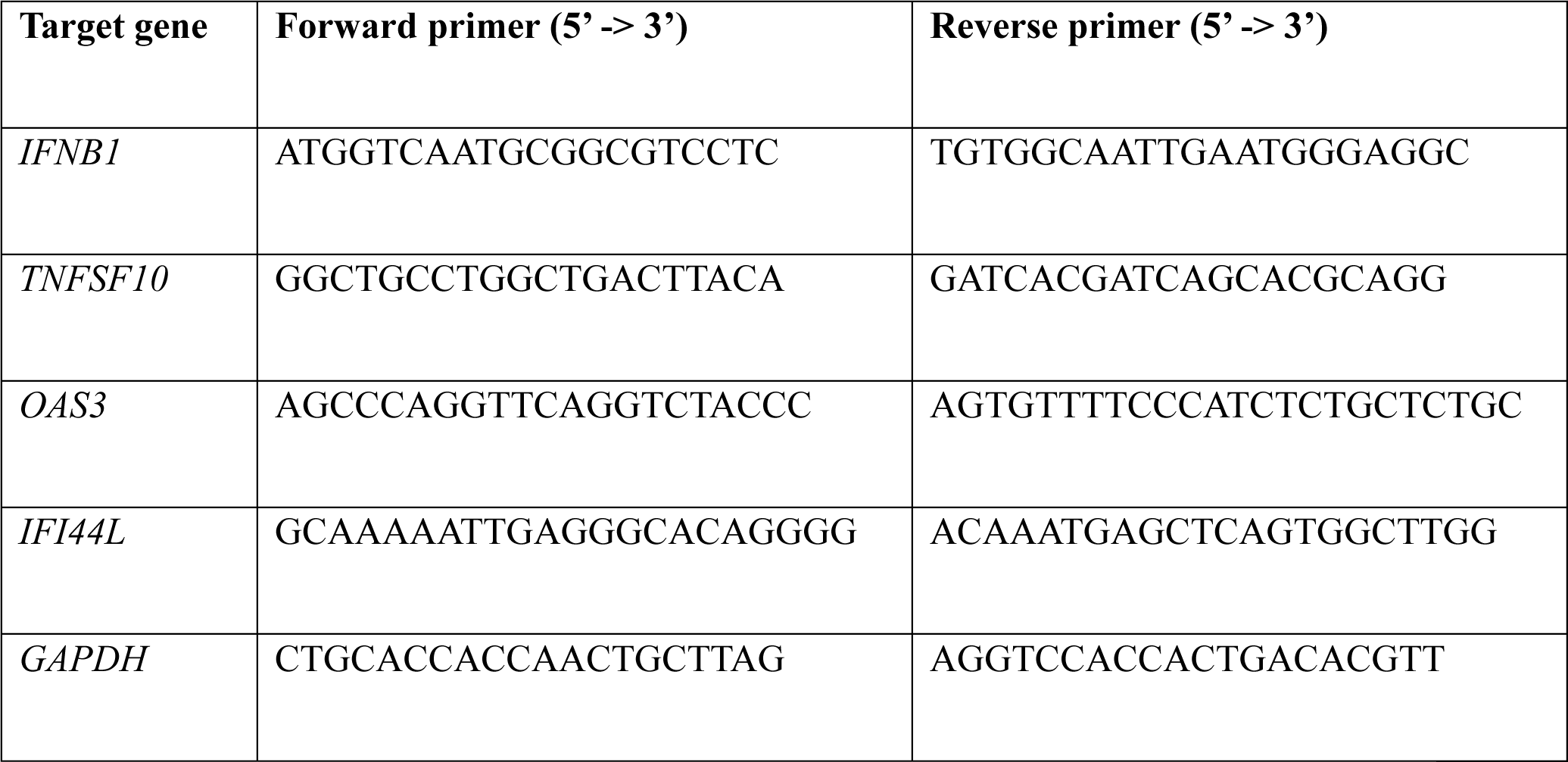
Primers sequences used for RT-qPCR.

### Western blot

Purified NK cells were treated with 100 nM TAK-981 for indicated time, equal number of cells were lysed with Laemmli buffer and run on SDS-PAGE. Antibodies against SUMO-1 (21C7), and SUMO2/3 (8A2) were obtained from the Developmental Studies Hybridoma Bank.

### Statistical analysis

Statistical analyses of the differences between data sets were performed using paired one-way ANOVA test, except for indicated experiment where paired student t-test was used (GraphPad Prism, GraphPad 10.0.2). Overall mice survival was estimated for each treatment group using the Kaplan-Meier method and compared with the log-rank test. P-values of less than 0.05 were considered significant (*, P < 0.05; **, P < 0.01; and ***, P < 0.001, ns = not significant).

## Authors contributions

RH designed and performed most of the experimental work. MdT conceived and performed in vivo experiments. DT analyzed RNA-Seq experiments and performed GSEA analysis. LC, SZ, DG, MV prepared the eNK. LG, SB, GC provided patient samples. SG performed the sequencing of RNA-Seq. MP provided IFNAR KO mice and tools. GB supervised the study and provided fundings. RH and GB wrote the manuscript. All authors edited the manuscript.

## Acknowledgments

We thank the members of the “Ubiquitin Family in Hematological Malignancies” group, Takeda Development Center Americas (Lexington, MA), for providing TAK-981. Funding was provided by the CNRS, the Fondation pour la Recherche Médicale (contract FDT202304016498 to RH; FDM201906008566 to LG), Institut National du Cancer-INCa_16072, the Fondation ARC pour la recherche sur le cancer. The HEMODIAG_2020 collection was funded by the Montpellier University Hospital, the Montpellier SIRIC and the Languedoc-Roussillon Region. MGX acknowledges financial support from the France Génomique National infrastructure, funded as part of “Investissements d’Avenir” program managed by the Agence Nationale pour la Recherche (contract ANR-10-INBS-09).

## References

1. Chretien AS, et al. High-dimensional mass cytometry analysis of NK cell alterations in AML identifies a subgroup with adverse clinical outcome. Proceedings of the National Academy of Sciences of the United States of America. 2021;118(22). 10.1073/pnas.2020459118.

2. D’Silva SZ, Singh M, Pinto AS. NK cell defects: implication in acute myeloid leukemia. Frontiers in Immunology. 2023;14. https://www.frontiersin.org/articles/10.3389/fimmu.2023.1112059. Accessed June 15, 2023.

3. Costello RT, et al. Defective expression and function of natural killer cell–triggering receptors in patients with acute myeloid leukemia. Blood. 2002;99(10):3661–3667.

4. Lion E, et al. Natural killer cell immune escape in acute myeloid leukemia. Leukemia 2012 26:9. 2012;26(9):2019–2026.

5. Xu J, Niu T. Natural killer cell-based immunotherapy for acute myeloid leukemia. Journal of Hematology & Oncology. 2020;13(1):167.

6. Boulanger M, et al. SUMO and Transcriptional Regulation: The Lessons of Large-Scale Proteomic, Modifomic and Genomic Studies. Molecules. 2021;26(4):828.

7. Boulanger M, et al. DeSUMOylation of chromatin-bound proteins limits the rapid transcriptional reprogramming induced by daunorubicin in acute myeloid leukemias. Nucleic Acids Research. 2023;gkad581.

8. Gabellier L, et al. SUMOylation inhibitor TAK-981 (subasumstat) synergizes with 5-azacytidine in preclinical models of acute myeloid leukemia. Haematologica. 2024;109(1):98–114.

9. Zitti B, et al. Innate immune activating ligand SUMOylation affects tumor cell recognition by NK cells. Scientific Reports. 2017;7. 10.1038/s41598-017-10403-0.

10. Kumar S, et al. Targeting pancreatic cancer by TAK-981: a SUMOylation inhibitor that activates the immune system and blocks cancer cell cycle progression in a preclinical model. Gut. 2022;71(11):2266–2283.

11. Nakamura A, et al. SUMOylation inhibitor subasumstat potentiates rituximab activity by IFN1-dependent macrophage and NK cell stimulation. Blood. 2022;139(18):2770–2781.

12. Lightcap ES, et al. A small-molecule SUMOylation inhibitor activates antitumor immune responses and potentiates immune therapies in preclinical models. Science Translational Medicine. 2021;13(611):eaba7791.

13. Demel UM, et al. Small molecule SUMO inhibition for biomarker-informed B-cell lymphoma therapy. Haematologica. 2023;108(2):555–567.

14. Lam V, et al. T cell-intrinsic immunomodulatory effects of TAK-981 (subasumstat), a SUMO-activating enzyme inhibitor, in chronic lymphocytic leukemia. Molecular Cancer Therapeutics. 2023;MCT-22-0762.

15. Dolstra H, et al. Successful Transfer of Umbilical Cord Blood CD34+ Hematopoietic Stem and Progenitor-derived NK Cells in Older Acute Myeloid Leukemia Patients. Clinical Cancer Research. 2017;23(15):4107–4118.

16. Wang S, et al. Breaking boundaries: Current progress of anticancer NK cell-based drug development. Drug Discovery Today. 2023;28(2):103436.

17. Crowl JT, Stetson DB. SUMO2 and SUMO3 redundantly prevent a noncanonical type I interferon response. PNAS. 2018;115(26):6798–6803.

18. Decque A, et al. Sumoylation coordinates the repression of inflammatory and anti-viral gene-expression programs during innate sensing. Nat Immunol. 2016;17(2):140–149.

19. Du L, et al. Mechanism of SUMOylation-Mediated Regulation of Type I IFN Expression. Journal of Molecular Biology. 2023;167968.

20. Nakagawa K, Yokosawa H. PIAS3 induces SUMO-1 modification and transcriptional repression of IRF-1. FEBS Letters. 2002;530(1–3):204–208.

21. Park J, et al. Elevated level of SUMOylated IRF-1 in tumor cells interferes with IRF-1-mediated apoptosis. PNAS. 2007;104(43):17028–17033.

22. Park S-M, et al. SUMOylated IRF-1 shows oncogenic potential by mimicking IRF-2. Biochemical and Biophysical Research Communications. 2010;391(1):926–930.

23. Kubota T, et al. Virus Infection Triggers SUMOylation of IRF3 and IRF7, Leading to the Negative Regulation of Type I Interferon Gene Expression*. Journal of Biological Chemistry. 2008;283(37):25660–25670.

24. Han K-J, Jiang L, Shu H-B. Regulation of IRF2 transcriptional activity by its sumoylation. Biochemical and Biophysical Research Communications. 2008;372(4):772–778.

25. Fu J, et al. MDA5 is SUMOylated by PIAS2β in the upregulation of Type I interferon signaling. Molecular Immunology. 2011;48(4):415–422.

26. Mi Z, et al. SUMOylation of RIG-I positively regulates the type I interferon signaling. Protein & Cell. 2010;1(3):275.

27. Hu M-M, et al. Innate immunity to RNA virus is regulated by temporal and reversible sumoylation of RIG-I and MDA5. Journal of Experimental Medicine. 2017;214(4):973–989.

28. Choi GW, et al. Formation of SUMO3-conjugated chains of MAVS induced by poly(dA:dT), a ligand of RIG-I, enhances the aggregation of MAVS that drives the secretion of interferon-β in human keratinocytes. Biochemical and Biophysical Research Communications. 2020;522(4):939–944.

29. Demel UM, et al. Activated SUMOylation restricts MHC class I antigen presentation to confer immune evasion in cancer. J Clin Invest. 2022;132(9):e152383.

30. Rey J, et al. Kinetics of Cytotoxic Lymphocytes Reconstitution after Induction Chemotherapy in Elderly AML Patients Reveals Progressive Recovery of Normal Phenotypic and Functional Features in NK Cells. Front Immunol. 2017;8:64.

31. Kim HS, et al. TAK-981, a SUMOylation inhibitor, suppresses AML growth immune-independently. Blood Advances. 2023;7(13):3155–3168.

32. Reina-Ortiz C, et al. Expanded NK cells from umbilical cord blood and adult peripheral blood combined with daratumumab are effective against tumor cells from multiple myeloma patients. OncoImmunology. 2021;10(1):1853314.

33. Sanchez-Martinez D, et al. Expansion Of Allogeneic NK Cells With Efficient Antibody-Dependent Cell Cytotoxicity Against Multiple Tumor Cells. Theranostic. 2018;8(14):3856–3869.

34. Kim D, et al. TopHat2: accurate alignment of transcriptomes in the presence of insertions, deletions and gene fusions. Genome Biol. 2013;14(4):R36.

35. Langmead B, Salzberg SL. Fast gapped-read alignment with Bowtie 2. Nat Methods. 2012;9(4):357–359.

36. Anders S, Pyl PT, Huber W. HTSeq—a Python framework to work with high-throughput sequencing data. Bioinformatics. 2015;31(2):166–169.

37. Love MI, Huber W, Anders S. Moderated estimation of fold change and dispersion for RNA-seq data with DESeq2. Genome Biol. 2014;15(12). 10.1186/s13059-014-0550-8.

38. Subramanian A, et al. Gene set enrichment analysis: A knowledge-based approach for interpreting genome-wide expression profiles. PNAS. 2005;102(43):15545–15550.

